# The spatio-temporal axis for phenotypic change: a comparison of source and translocated Arctic charr populations after 25 generations

**DOI:** 10.1101/2020.12.08.416073

**Authors:** Marius H. Hassve, Mari Hagenlund, Kjartan Østbye, Katja Häkli, Thomas Vogler, Finn Gregersen, Gørli B. Andersen, Svein O. Stegarud, Kjell Langdal, Max E. Waalberg, Karl C. Langevoll, Kim Præbel

## Abstract

Evolution of morphological traits is hypothesized to act on an extended time scale, yet studies have suggested that these changes are possible within a few generations. Trophic polymorphism enabled through niche adaptations and ecological opportunity is one phenomenon that facilitate occurrence of rapid adaptive variation, common in many northern freshwater fish species. One such species is Arctic charr, which is known for its extensive variation in morphology and the occurrence of morphs. However, the speed at which such morphological variation arises is poorly studied despite the importance for understanding the onset of evolution. The aim of this study was to elucidate this process in a gradient of eight lakes that was stocked with Arctic charr in the period from 1910 to 1917 from Lake Tinnsjøen, Norway. We used morphological measurements to test for differences in traits between populations and Haldane and Darwin’s evolutionary rates to estimate divergence rates in traits. We also tested for correlation between putative genetic and morphological divergence. In addition, we contrasted the morphological divergence with that expected under neutral genetic expectations, using 12 microsatellite markers, to analyze whether and which morphological differences that is following early genetic divergence. A significant genetic differentiation was found between the source population and five of the translocated populations with corresponding differences in morphological traits for four of the populations. Population genetic structuring indicated six different genetic clusters. The translocated populations also exhibited trait divergence estimated with both Haldane and Darwin’s rates. Differences in morphological traits showed a significant correlation with genetic divergence, and the morphological differences were most likely affected by differences in lake parameters such as maximum depth, lake size and fish community. We conclude that intraspecific morphological and genetic divergence can form on short evolutionary time scales with important implications for conservation and management practices.

## Introduction

Adaptive changes in phenotypic traits mediated by natural selection is a crucial component in evolution resulting in population differentiation (1). In this process, adaptation to different physical environments, such as divergent niches, temperature and climate regimes, as well as intra- and interspecific interactions may be important factors (2–4). The evolution of morphological trait divergence is assumed to act on a long time scale, however this ongoing process can also act on a contemporary time scale even within a few generations (5, 6). For example, the body and beak size of the birds *Geospiza fortis* and *G. scandens* on the Galapagos Islands have changed within 30 years due to environmental selection pressures exerted by El Nino climate conditions (7). Other examples of short time-based trait-divergences among populations within a set of different taxonomic groups imply that rapid divergence and contemporary evolution may be a common feature in nature (6, 8–10).

A phenomenon that suggests early ongoing adaptive divergence within species is the presence of sympatric trophic polymorphism, which is frequently observed within some of the northern freshwater fish taxa such as *Salvelinus*, *Coregonus* and *Gasterosteus* (11–15). Such trophic polymorphism may result from adaptive diversification of morphological traits driven by heterogeneous resource utilization, which could lead to specialization to specific niches given ecological opportunity and sufficient selection pressures (16, 17). The availability of divergent lake niches seems to be determined by both abiotic and biotic variables such as temperature, lake morphometry and biotic interactions (18, 19). Fennoscandian freshwater lakes are often found to harbour few fish species (20), potentially reducing the level of interspecific competition. Thus, contingent upon selection pressure, the evolution of trophic polymorphism in the ecological speciation process will likely vary in strength in different environments. The ecological speciation process is important in the rapid formation of new species through adaptive radiation (17).

Arctic charr (*Salvelinus alpinus* L.) is a circumpolar, coldwater adapted, and highly polymorphic species (21–23), known for its extensive variation in morphology both among and within lakes (24–26). One to four morphological variants (further referred to as morphs of Arctic charr may co-occur in the same lake but show low to moderate genetic differentiation among sympatric morphs (15, 22, 24, 27). This diversity of morphological variants gave rise to what has been termed “*the charr problem”* which still puzzles scientists with regard to whether the formation of sympatric morphs have an allopatric or sympatric origin (22, 28, 29). Due to this complexity, there seem to exist taxonomic confusion with regard to assigning Latin names to Arctic charr populations and morphs within the “*charr complex”* (30–32). Studies considering the evolutionary origin of morphs (33), taxonomy (34–36), and phylogeny (33, 37, 38) are common in the literature. According to Kottelat & Freyhof’s (32) taxonomic evaluation, there exists over 30 species of the genus *Salvelinus* in Europe. Their system, which may challenge general acceptance in the scientific community, is supported by the IUCN Freshwater Biodiversity Unit (32), potentially creating precedence for future management of the Arctic charr biodiversity. However, regardless of whether the Arctic charr complex consists of different species or not, conserving biodiversity and genetic heterogeneity is of high importance where examples of management of species at population/unit level has been implemented to conserve uniqueness (39–41). Although such evaluations are now common in e.g., North America (42), currently it raises little management focus in most of European countries.

Monomorphic and polymorphic Arctic charr populations may vary in external and internal characteristics (32, 43, 44), including morphometric and meristic characters such as e.g., body and head proportions (45–49), number and length of gill rakers (28, 35, 50, 51), and length of the pectoral fin (26). Trait variation is often associated with niche utilization (51, 52), as seen in other postglacial fishes (e.g. 12). Here, two general patterns are seen where planktivorous Arctic charr morphs have many gill rakers (53, 54) which is important for feeding in the pelagic zone (51), and the benthivorous morphs have large pectoral fins for efficient maneuvering in the littoral niche (45). These two traits have been suggested to show parallel evolution within the Arctic charr complex (55). Crossing experiments in common garden facilities show that these traits may have a genetic basis (22). Few have estimated the rate of differentiation or adaptation in phenotypic traits in Arctic charr. Michaud, Power (56) found divergence in fins, maxilla, and growth in Arctic charr after six generations following introduction of charr into a new system, implying rapid changes. Support for rapid change is also found in sockeye salmon (*Oncorhynchus nerka*) which diversified into two reproductive isolated populations after 13 generations following introduction, resulting in significant differences in body shape (57).

In Lake Tinnsjøen, Norway, the Arctic charr have been documented to consist of four morphologically and genetically distinct morphs; the planktivore, dwarf, piscivore-, and abyssal morph (58). The Arctic charr in Lake Tinnsjøen have historically acted as a source population for large scale translocation of hatchery reared offspring of wild parents (59 [Fisheries Inspector]). It was most likely the planktivorous Arctic charr that were used for stocking/translocation, as this morph is considered the most valuable for food among the morphs. From the year 1900 to 1915, stocking of freshwater fish was a recommended action from the Norwegian government (60). Hence, out of Norway’s more than 30,000 Arctic charr populations (61), it is likely that many populations are stocked and do not originate from natural immigration after the last glaciation (20). Populations where documentation for spatio-temporal translocation exist offers an opportunity for testing hypotheses related to if and how divergence at morphological traits and genetic loci, and the interaction between genetic and morphological divergence contribute to formation of new morphs within species. Genetic diversity of founders may determine the scope for tackling environmental change (62), due to its basis for phenotypic diversification. Divergent traits among populations may also reflect phenotypic plasticity and genetic drift in small founder populations (63). Changes in morphology may be due to selection on heritable traits (64, 65), where also phenotypic plasticity (66), may play a role when comparing source- and translocated freshwater fish populations within a time interval of 100 years of isolation (67).

The Lake Tinnsjøen Arctic charr were translocated approximately 100 years ago (between 1910-1917) into eight lakes and, if assuming a generation length of four years (58), the stocked populations have evolved in allopatry for approximately 25 generations. Hence, the system offers excellent opportunity for illuminating the questions of if, and how fast, traits diversify, and gene pools diverge in different environments. This is a key point when managing or conserving populations, morphs, or species, especially in light of predicted climate scenarios and habitat loss. This study encompasses a large-scale natural experiment that investigate founder effects and varying responses in Arctic charr populations translocated from a source population to more or less replicated spatially separated environments 100 years ago, lending a novel opportunity to study phenotypic and genetic changes. Thus, the aim of the study was to investigate the morphological and genetic response as well as rates of differentiation by studying translocations of Lake Tinnsjøen Arctic charr into neighboring, but different, lake environments.

## Materials and methods

### Study systems

A hatchery existed in Lake Tinnsjøen producing offspring of wild caught Arctic charr parents where the main intention was to stock Arctic charr in nearby lakes for human consumption. We assume that the most common Arctic charr morph (which is also the most appreciated as a food source) was used for breeding in the hatchery. This is the planktivorous morph (referred to as the “normal morph” or “pelagic charr“). More details of the diversity of the Lake Tinnsjøen Arctic charr morphs are given in (58). The study design contrasted the source population in Lake Tinnsjøen (TI) with eight translocated populations; Lake Torvevatnet (TO), Tjyrutjønn (TJ), Sønstevatn (SØ), Finsevatn (FI), Lufsjå (LU), Store Snellåtjønn (SN), Grautnattjønnan (GR) and Svømhovdtjønn (SV) (Table 1, Fig. 1). The selection of lakes was based on information from description of 50 lakes with known introduction history (59). The year of stocking and the number of translocated Arctic charr juveniles are specified in (59; Table 1). With regard to Lake Finsevatn, the introduction probably took place in connection with the introduction of 30 000 charr in surrounding lakes at this time (68).

**Table 1.** Morphometric and water chemical description of lakes where Arctic charr were sampled with sampling dates, number (N) of Arctic charr samples, fish species present, year of release, and number of charr released in-or in areas surrounding the Lakes. Sampling date was as follows: FI - August 2012; TJ, GR and SV - July 2014; TO - September 2014; LU and SØ – September and October – 2014; SN – August 2014 and February 2015; TI – July and October 2013.

| Lake | M.a.s.l. | Surface<br>area<br>(km<sup>2</sup>) | Max<br>depth<br>(m) | Secchi<br>depth<br>(m) | pH<sup>#</sup> | A<sup>#</sup> | C<sup>#</sup> | T.H<sup>#</sup> | N<br>cha<br>rr | Fish<br>species* | Year<br>stocked | Number<br>stocked |
| --- | --- | --- | --- | --- | --- | --- | --- | --- | --- | --- | --- | --- |
| TI | 191 | 51.5 | 460 | 9 | 6.55 | 110 | 14.18 | 168 | 28 | Sa, St,<br>Pf, Pp | --- | ----- |
| TJ | 1068 | 0.04 | 2 | 2 | 6.73 | 40 | 9.17 | 110 | 25 | Sa, St | 1913 | 1000 |
| GR | 1069 | 0.09 | 35 | 12 | 6.28 | 14 | 6.23 | 88 | 29 | Sa | 1917 | 1000 |
| SV | 1080 | 0.1 | 10 | 10 | 6.10 | 10 | 5.98 | 100 | 26 | Sa, St | 1917 | 1000 |
| LU | 1214 | 4.9 | 10 | 10 | 6.72 | 34 | 8.74 | 100 | 29 | Sa, St | 1913 | 10 000 |
| SN | 1176 | 0.1 | 12 | 8 | 6.59 | 34 | 8.12 | 90 | 13 | Sa, St | 1916 | 2000 |
| TO | 1025 | 0.8 | 38 | 6 | 6.87 | 44 | 10.34 | 106 | 24 | Sa, St | 1913 | 3000 |
| SØ | 317 | 2.5 | 30 | 5 | 7.34 | 78 | 21.30 | 190 | 30 | Sa, St,<br>Pf, Cc | 1914 | 1500 |
| FI<sup>#</sup> | 1216 | 3.2 | 32 | 6 | 6.89 | 81 | 17.70 | 124 | 25 | Sa | 1910 | 30 000 |

\*Fish species codes; Sa = *Salvelinus alpinus*, St = *Salmo trutta*, Pf = *Perca fluviatilis*, Cc= *Carassius carassius*, and Pp = *Phoxinus phoxinus*. # Lake morphometry are retrieved from Waalberg (68) and water chemistry information from Aanes (69). T.H=Total hardness (µekv/l), C=Conductivity (µS/cm), A=Alkalinity (µekv/l).

**Fig. 1.**
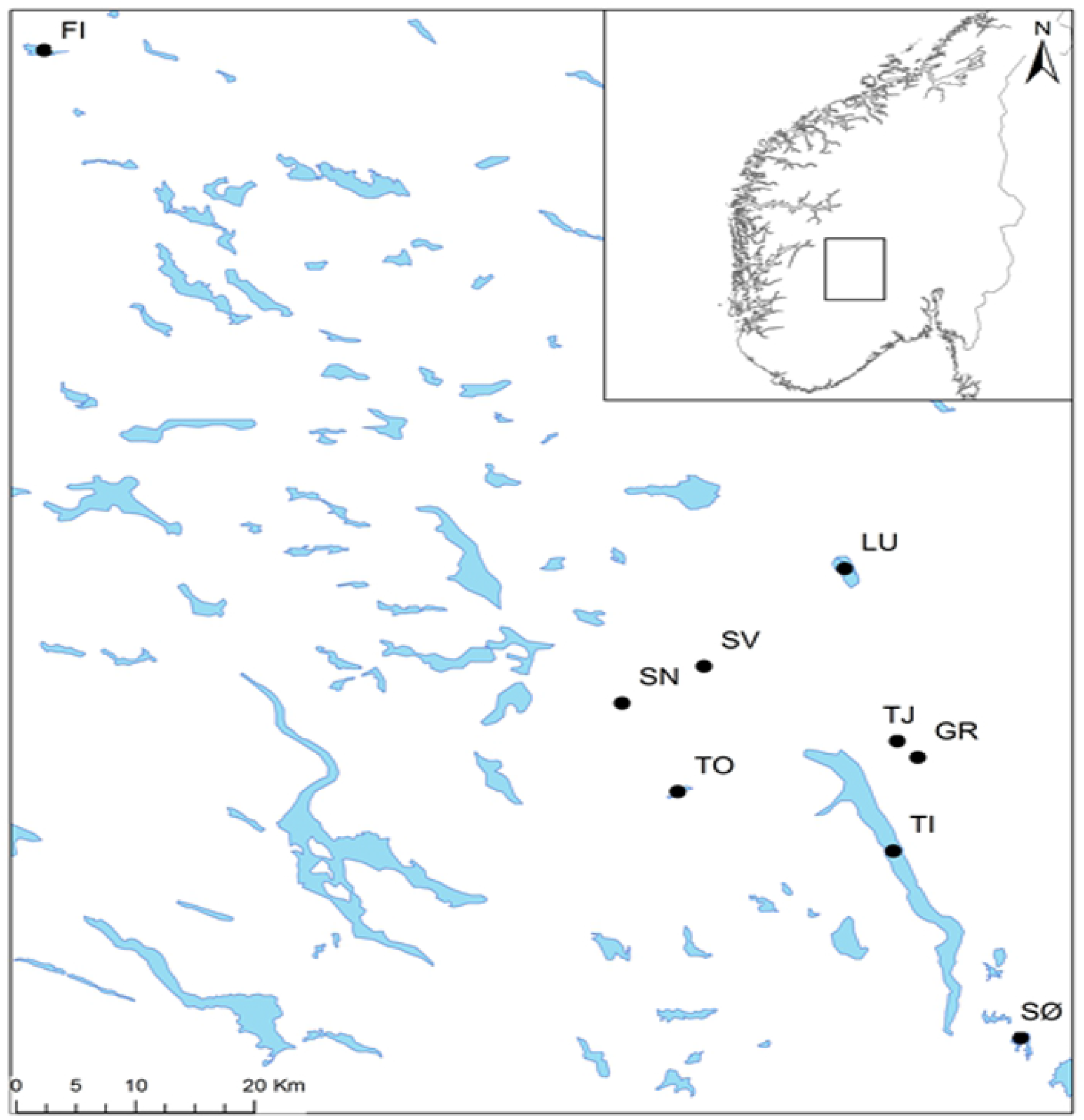
Map over the source Lake Tinnsjøen (TI) and the translocated populations, with corresponding overview where the lakes are centered geographically in southern Norway.

All the translocated populations, except Lake Finsevatn (FI), are geographically in close proximity to the source population of Lake Tinnsjøen (TI), in Tinn and Notodden municipality, Telemark County, Norway (Fig. 1). All lakes varied in elevation, surface area and maximum depth, with Lake TI having the lowest elevation, largest surface area, and deepest maximum depth (Table 1). Temperature was measured for one year, from July 2014 to July 2015, at 4 meters depth with an iBCod 22L temperature logger in a shallow (TJ), medium (SV) - and deep (GR) lake that were stocked with charr in the mountain areas. For TI only temperatures from June are available from literature measured at depths ranging from 10-20 meters (70).

### Sample collection

For both the source population in Lake TI and the specific translocated population of Lake FI a representative subset of Arctic charr from the littoral, pelagic and profundal zones were selected from a larger sample and used for further analyses. The sampling effort and datasets are fully described in Langevoll (71), Waalberg (68) and (58). For TI specifically, only individuals from the planktivorous Arctic charr morph were contrasted to the other lakes. The selection of the planktivorous Arctic charr individuals from Lake TI were based on a geometric landmark analysis described and conducted and in (58).

Gill-nets (Nordic multimesh gillnet series) (mesh size described in (58)) were set in all lake habitats (littoral, pelagial and profundal). Benthic gillnets were placed from land in the littoral zone, while profundal gill-nets were placed at the deepest point in each lake, and floating gillnets were placed in the open water at depths between 0-6 meters. In accordance with lake size, depth, species composition, and local knowledge, catch effort was adjusted among localities from a baseline of four littoral- and profundal nets, and one pelagic net. In a medium sized lake such as LU, we increased the effort in the pelagic area to three gillnets, while in the smallest lake, TJ, only four nets were set in total. All nets were set to fish overnight for 12-hours.

After catch, the right pectoral fin from all Arctic charr was stored on 95% ethanol for DNA analysis. Fish were stored in an ice cooler and brought to a freezer (−40 °C) for storage. The length in mm (from the tip of the snout to the end of the caudal fin) was measured in all fish.

From all lakes (except FI) we measured transparency and water color using a secchi disc, while maximum depth was recorded using a handheld echo sounder (Plastimo Echotest II). Water samples were taken from the surface level at approximately 20 cm depth in the pelagic area (approximately equal distance to shoreline from all angles) and stored in a freezer in dark containers for later analysis of pH, conductivity, alkalinity and total hardness. To obtain a rough estimate of the invertebrate community (potential prey items) in the different lakes we used a standardized kick-sample procedure in the littoral area with an associated dip net. Zooplankton was sampled in the pelagic zone using a sieve of 250 μm mesh size from around 20 meters depth up to the surface vertically (in shallow lakes from right above the lakebed). Unfortunately, no sampling of invertebrate or zooplankton was performed in Lake FI.

### Morphological analyses

Photos from 227 Arctic charr were taken (after freezing) using a Canon EOS 550d camera and Canon lens, EFS 18 – 55 mm with constant focal length and exposure settings (F20 ISO1600 AV with blitz). Each individual where photographed lying in a natural extended position with the left side up, using a standardized light setting (CL – Led 15). Photos were then digitized into tps files in tpsUtil version 1.53 (72), before being transferred to tpsDig2 version 2.16 (73), were a set of the 30 fixed and sliding label landmarks used in (58) were set (Fig. 2). In MorphoJ (74), extreme outliers were removed from the dataset in accordance with the results from an outlier analysis followed by a visual inspection of the individual fish in question and a Procrustes fit analysis. A principal component analysis (PCA) with corresponding eigenvalues was calculated for all landmarks to describe differences in individual variation in body shape. To correct for length effects a regression between shape data and log centroid size was performed in MorphoJ (74), saving the residuals. A PCA analysis was performed in MorphoJ using residuals. The PC scores were exported for further analyses.

**Fig. 2.**
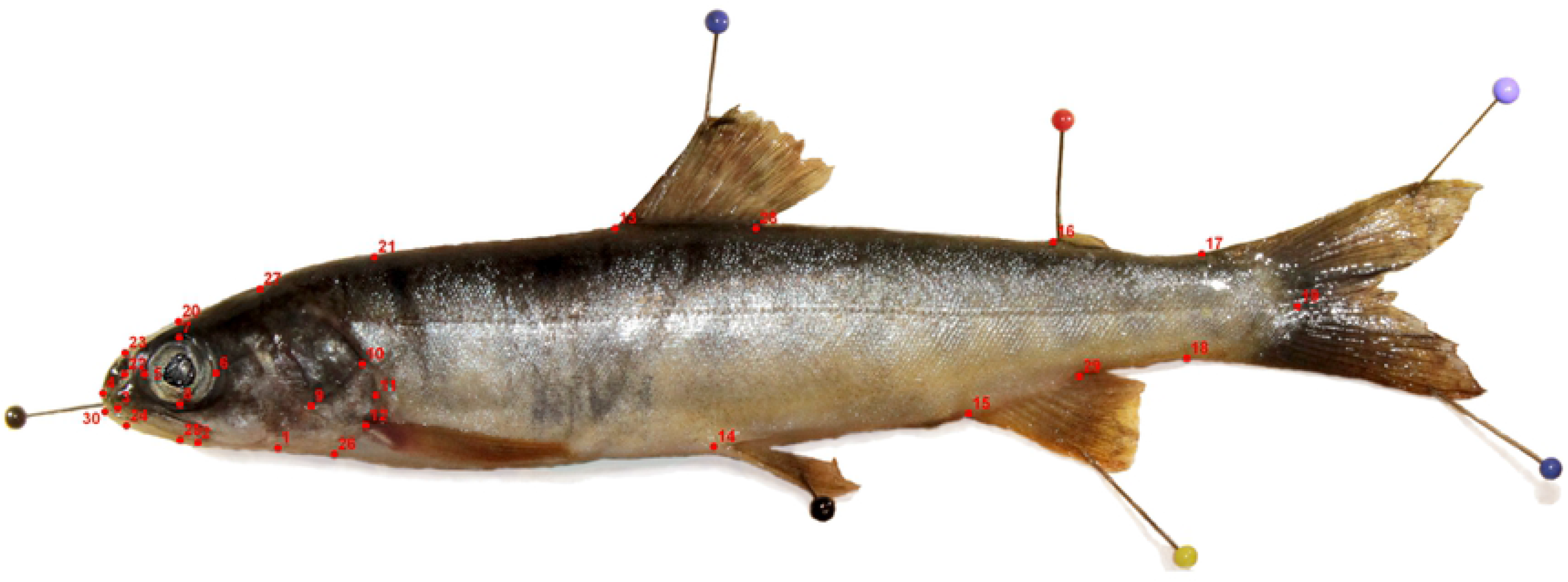
Detailed explanations of landmark positions: 1;lower edge of preoperculum, 2;edge of maxillary bone, 3;mouth opening, 4; tip of the snout, 5 – 8; eye positions, 9; mid edge of preoperculum, 10;posterior edge of preoperculum, 11; posterior edge of operculum, 12; pectoral fin, 13 and 28; dorsal fin, 14; pelvic fin, 15 and 29; anal fin, 16; adipose fin, 17; upper tail root, 18; lower tail root, 19; end of the side line organ, 20; top of head, 21; back above pectoral fin, 22; nostril, 23; over nostril, 24; under-jaw, 25; edge of mouth, 26; lower edge of operculum, 27; transition zone from head to body, 30; edge of lower lip.

In Mclust 5.01 (75) we tested how many normal distributions that were present using model-based clustering and visualization based on Bayesian Information Criterion (BIC) when analyzing the three first PC axes. PC 1 explained 20.88 %, PC 2 19.41 %, and PC 3 14.08 %, giving a total explained variance of 53.95 % (Fig.S1). Furthermore, all the 9 default models in Mclust were tested using these three PC axes. The model with the highest BIC value was then selected as the most parsimonious model. In cases where BIC values had less than 2 units in difference, models were considered as equal.

A set of eight metric and two meristic traits were analyzed as these traits may reflect phenotypic plasticity or adaptations to different environments. First, the length of the left pectoral fin (LP), upper maxilla length (M), head length (HL), and eye-area (EA (***πr***^2^)) was measured using a Mitutoyo 500-606 caliper (0.1 mm precision). Secondly, the first left gill arch was dissected out and the lamella length (L) and the length of the second gill raker (in the angle between the upper and lower gill arch (LG)) were measured using a Leica microscope (ocular 16x/14B) with an internal ruler (measurements converted to real size in mm). The length of the upper (UA-L) and lower gill arch (LA-L) were measured using a digital caliper. The number of gill rakers on the upper (GR-U) and lower gill arch (GR-L) was counted in the microscope.

For eight metric and meristic traits (GR-U, GR-L, LG, M, L, LH, EA and LP), univariate ANOVA`s were performed between the nine lakes to test if there were statistical differences between traits in the nine lakes. This analysis was performed after correcting for fish body size using residuals from regressions singularly of length versus all the traits in R 3.1.3 (76).

A multivariate linear discriminant analysis with all 10 traits combined were conducted in the Ade4 package in R 3.1.3 (77) using residuals, calculating the most likely number of phenotypic clusters across the nine lakes. In addition, all the individuals were subjected to a “back-assignment” procedure where individuals are mixed independently of original population before being assigned to populations by using all the morphological traits.

A multivariate analysis of variance (MANOVA) was performed in R 3.1.3 (76) using morphological and meristic traits from the 9 populations, together with maximum depth, lake size (km), presence of multi-or single fish species communities, and individual lakes. The dependent variables were; M, LP, LH, GR-U, L, GR-L, LG, LA-L, EA, UA-L, PC1 and PC2. PC1 and PC2 are the first two PC scores from the PCA analysis in MorphoJ describing the body shape of the fish. Only the first two PC axes were used for this analysis as they explained approximately 40 % of the variance. Meters above sea level (MASL) was not included in the analysis since seven of the nine lakes are located at approximately the same altitude.

### Estimating *Haldane’s* and *Darwin’s* rates

The divergence rates for all the ten phenotypic traits between the source population of Lake Tinnsjøen and the eight translocated populations were estimated using *Haldane’s* and *Darwin’s* (5). Here, the time unit (*t*) for the Darwin analysis was set to 100 years, corresponding to 25 generations (g) for the Haldane analysis. The generation time was estimated to be 4 years based on the youngest mature planktivorous Arctic charr found in Lake Tinnsjøen. This estimated generation time is based on the methods used by Michaud, Power (56) who studied translocated Arctic charr. We further estimated the elapsed number of generations between source and translocated populations to be approximately 25 generations.

### Genetic analysis

DNA was extracted from the pectoral fin using the *Omega Biotek E-Z 96 Tissue DNA Kit* for the translocated populations (2015), and *QIAGEN DNeasy Blood & Tissue Kit* for the source population (2013), following the manufactures protocols, with some minor adjustments. The DNA concentration were estimated on a representative subset of all the samples by nano-drop (ND – 1000) to 89.1-189.9 ng/uL and 77.5-370.8 ng/uL, E-Z 96 and DNeasy, respectively. 12 microsatellite loci were amplified by polymerase chain reaction (PCR) in two multiplex panels. Multiplex one consisted of the primers; *OMM1105*, *SCO220*, *SALP61SFU*, *SCO212*, *SALF56SFU*, and *SCO218*, while multiplex two consisted of *SMA17*, *SFO334LAV*, *SCO204*, *SALJ81SFU*, *SCO215*, and *SMA22*. Amplifications were performed in 2.5 μl PCR reactions with 5-10 ng template DNA, 1.25 μl Qiagen multiplex master mix, 0.25 μl primer mix (multiplex 1 & 2), and 0.5 μl ddH_2_O, using PCR conditions in (78). The PCR products were analysed on an ABI 3130 XL Automated Genetic Analyzer (Applied Biosystems) and the alleles were scored using the GENEMAPPER 3.7 software (Applied Biosystems). All scores were visually inspected to verify the allele binning, blank and replicate samples. No contamination or miss-matches between replicates were observed.

The locus *SCO215* was monomorphic and omitted from the dataset. To check for genotyping errors such as null-alleles, stutter errors, large allele drop-out, and size-independent allelic drop-out, we used Micro-checker 2.2.3 (79). Micro-checker revealed homozygote excess, possibly due to null alleles, in 3 of 12 loci; *SCO212* in population SV, *SFO334LAV* in populations SN, TI and TJ, and *SCO220* in populations LU, SØ and SV. Since loci *SFO334LAV* and *SCO220* showed homozygote excess in three populations, the program FREENA (80, 81) was run to check for null alleles. Null alleles may cause an overestimation of genetic differentiation (*F_ST_*) and distance, however FREENA corrects for allele-frequency biases by using the *Excluding Null Alleles* (ENA) method (80). FREENA was run with 5000 iterations, where the corrected and uncorrected *F_ST_* values were compared using a pairwise t-test that did not reveal a significant difference (Pairwise t-test: t_35_=1.2, p=0.23). Since the occurrence of homozygote excess did not seem to have a significant effect on *F_ST_*, it is highly unlikely that the presence of homozygote excess in some of the loci will have a substantial effect on subsequent genetic analyses in our study.

Loci that are under selection may affect genetic structure and genetic differentiation between populations (82). Thus, the software Lositan (82, 83) was used to test if any of the 12 loci were candidates for positive-or balancing selection. All loci were run under both the stepwise mutation model (SMM) and the infinite allele’s method (IAM) with the “Force mean F_ST_” and “Neutral mean F_ST_” alternatives. All analyses were run with 100000 simulations. Both the SMM and the IAM models indicated that loci *SFO334LAV* and *SCO220* were candidates for positive selection. Due to these findings and occurrence of homozygote excess in several populations for these two loci, loci *SFO334LAV* and *SCO220* were excluded from subsequent analyses. To check for deviations from Hardy-Weinberg (HWE) and linkage disequilibrium (LD), Genepop 4.2 (84) was run using exact tests. To further test for significant values of HWE and LD, the False Discovery Rate (FDR) corrections (85) were implemented to correct for false positives when conducting multiple tests. Out of the 234 tests, FDR indicated significant departures from HWE in six individuals. There was no evidence of a systematic departure from HWE in any of the populations (i.e. none of the significant markers were consequent throughout populations). Out of 405 tests, significant LD was found in 11 loci pairs after FDR correction, and 7 of these significant findings were systematic for the loci pair: *SCO218* and *SCO204*. As LD may influence genetic assignment, a test was performed where *SCO204* was removed and the hierarchical STRUCTURE approach (see below) was run again to test for differences in population assignment. No difference was found in any of the runs in Structure, and *SCO204* was thus included in subsequent analyses.

Genetic diversity estimates such as mean and standard error of the number of alleles (*N_a_*), number of effective alleles (*N_e_*), observed-(*H_o_*) and expected heterozygosity (*He*) and unbiased expected heterozygosity (*uHe*) were calculated in Genalex 6.5 (86), as well as the software Fstat 2.9.3. (87) (Table S1). An *F_ST_* matrix for all population pairs was calculated in FREENA (80), which calculates *F_ST_* with- and without the ENA method. P-values for significant differentiation in *F_ST_* values was calculated for each population pair and corrected for multiple tests with False Discovery Rate correction (85). Standardized private allelic richness (*Ap*) and standardized allelic richness (*Ar*) which corrects for differences in sample size was calculated using the software HP-rare (Table S2) 1.0 (88), using N=24 as the smallest sample size.

### Population genetic analyses

STRUCTURE 2.3.4 (89) was used to determine the most likely number of genetic clusters and to determine similarity in genetic structure between Lake Tinnsjøen and translocated populations. The admixture model was chosen for the initial analysis, including the LOCPRIOR function which considers geographic sampling information as recommended by (90). Structure was run 10 times for each K with 100,000 burn-ins followed by 100,000 Markov Chain Monte-Carlo repetitions. In addition, a hierarchical method was implemented to determine which populations grouped together as recommended by Evanno, Regnaut (91). The online program Structure Harvester (92) was used to determine the most likely number of clusters based on the values of both LnP(K) and deltaK (Fig. S2).

A phylogenetic neighbor-joining tree that groups populations based on NEI`s minimum genetic distance (93) was built in Populations 1.2.32 (94). 100 bootstrap repetitions was performed and visualized in TreeView32 (95). A second phylogenetic neighbor-joining tree, including five outgroups from southern and northern Norway was performed to evaluate clustering (Fig. S4).

### Association between genetic and phenotypic variation

To assess the genetic diversity among individuals and populations, a principal component approach was used to identify the axes of greatest genetic differentiation. A PCA of allele frequencies on microsatellite data was performed with Adegenet package in R (96), and then used in further analysis to illustrate the genetic differences between individuals.

Redundancy analysis (RDA) was performed to assess how much of the genetic variation reflected the observed phenotypic variation, and to detect the main traits that associated with the genotypic variation. In order to prevent overfitting, the command “OrdiStep” in the R package VEGAN, was used for forward selection of morphological traits using residuals (97).

Overall *F_ST_* was calculated with the Hierfstat R package (98). As the total variation of a PCA on allele frequencies correspond to *F_ST_* (99), it is possible to multiply the percentage of constrained variation in RDA by the overall value of *F_ST_* to obtain the equivalent *F_ST_* for proportion of the total genetic variation that is explained by the explanatory variables (100).

## Results

### Biotic and abiotic characteristics of the lakes

Two lowland and seven mountain lakes were sampled. The lake size varied from 0.04 (TJ) to 51.5 (TI) km^2^ and maximum depth varied from 2 (TJ) to 460 (TI) meters (Table 1). For Lake TI only temperatures from June are available from the literature, with an approximate temperature of 4.8°C measured at depths ranging from 10-20 meters (70). In comparison, the 35 meter deep Lake GR had the lowest mean temperature from June of 4.7°C (yearly temperature sum of 1437 °C), while the 10 meters deep Lake SV had mean June temperatures of 5.4°C (yearly temperature sum of 1929 °C) and the 2 meter deep Lake TJ had a mean June temperature of 7.9°C (yearly temperature sum of 2013 °C). The invertebrate and zooplankton communities were relatively similar in composition for both the lowland and mountain lakes, consisting of invertebrates such as chironomid larvae, and oligochaetes, as well as zooplankton such as *Daphnia longispina*, *Holopedium gibberum* and Copepods (Table S3). No calculations were performed on the invertebrate or zooplankton communities.

### Morphological analyses

The Mclust analysis for differences in body shape indicated two models with equal BIC values; both indicating one single cluster where the first had equal, spherical shape, and equal volume (EII, 1, BIC: 4453), while the second model (VII, 1, BIC: 4453) had variable volume (df = 4, n = 257, Fig. S3). According to the most parsimonious model in Mclust, the Arctic charr from the eight translocated lakes (FI, GR, LU, SN, SV, SØ, TJ, TO) and the source population (TI) clustered into one single group with regard to body shape (Fig. 3 & S3).

**Fig. 3.**
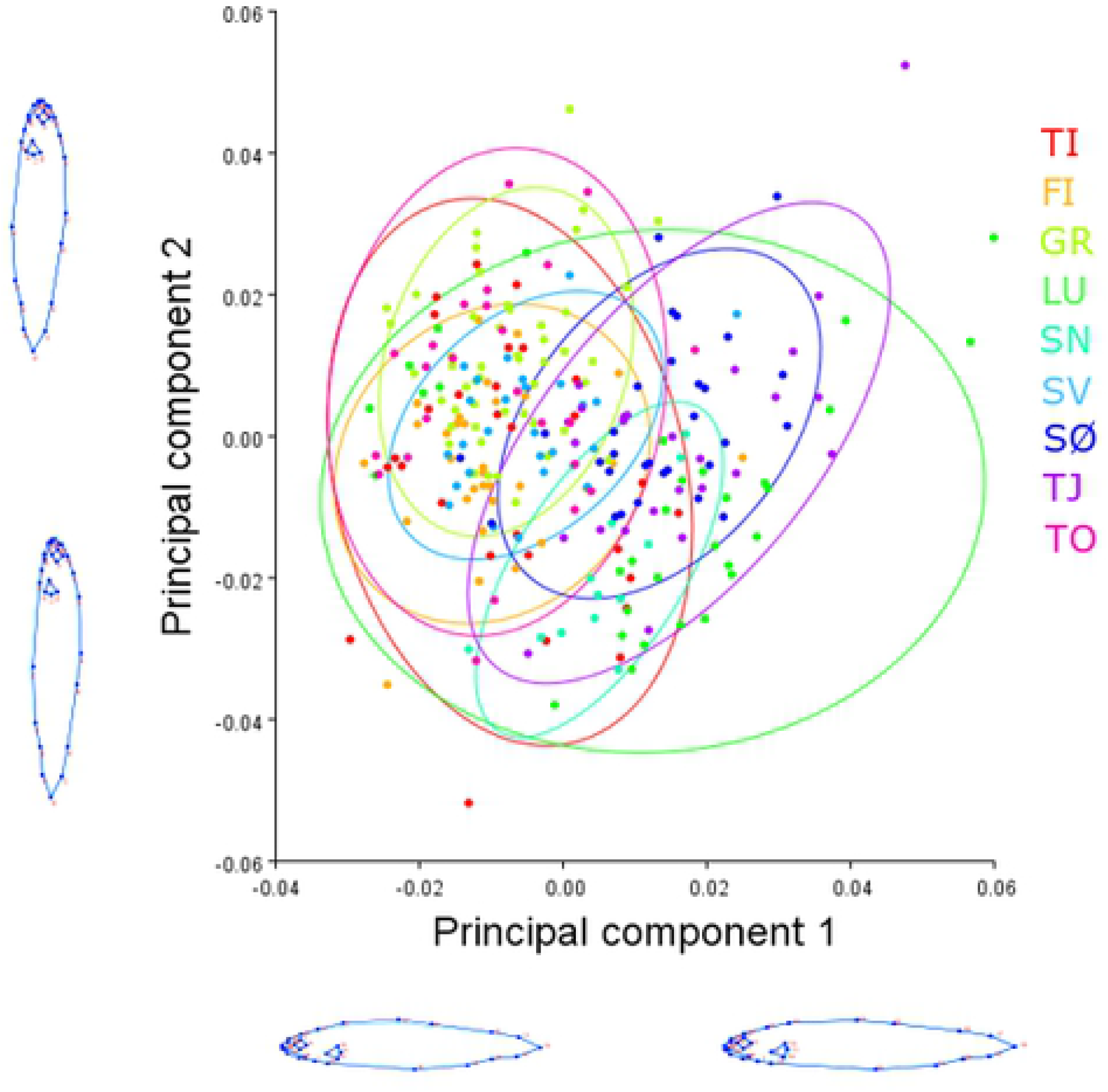
Morphological variation (landmark - shape) from source- and translocated Arctic charr populations. The different colors represent the source- and translocated populations. Population TI is marked as red circle, FI as yellow, GR as light-green, LU as dark-green, SN as turquoise, SV as light-blue, SØ as dark-blue, TJ as purple and TO as pink. Ellipses represent 95% confidence intervals.

Univariate ANOVA’s showed significant differences in metric and meristic traits among the translocated Arctic charr. Between the source and translocated populations, the only significant difference was found in the number of gill rakers on the lower raker (GR-L, Table 2), where Lake TI charr had to have the greatest number of GR-L (mean 15.7). Among the translocated populations, several traits differed significantly; the number of gill rakers on the upper arch (GR-U) differed significantly between Lake FI, LU, SV, TJ and TO with Lake SV having the highest number (mean 10.5) and Lake TJ having the lowest number (mean 9.6). The number of gill rakers on the lower arch (GR-L) differed between Lake FI, GR and TJ (in addition to TI), where Lake FI had the highest number (mean 15.2) and Lake GR had the lowest number (mean 14.8). For the longest gill raker (LG) significant differences were found between Arctic charr from; FI, GR, SN, SV, SØ, TJ and TO, where Lake TJ exhibited the longest LG (5.1), and Lake GR the shortest LG (3.0). The length of the mandibula (M) differed significantly between populations; FI, GR, SØ and TJ, where the longest mandibula was found in Lake TJ (mean 25.9), and the smallest in Lake GR (mean 12.5). For length of the lamella (L), all translocated Arctic charr exhibited significant differences, with Lake SN having the longest lamella (mean 7.2), and Lake GR exhibiting the shortest lamella (mean 3.2). Head length (LH) differed significantly for Arctic charr from Lake FI, GR, SV, SØ, TJ and TO. Arctic charr from Lake TJ exhibited the longest head length (mean 51.9), and Arctic charr from Lake GR had the shortest head length (mean 27.5). Eye area size (EA) differed significantly between Arctic charr from Lake GR and SV, where Arctic charr from Lake SV had the largest eye area (mean 44.5) and Arctic charr from Lake GR had the smallest eye area (mean 27.6). Finally, length of the pectoral fin differed significantly between Arctic charr from Lake GR, SV, SØ and TJ. Here, Arctic charr from Lake TJ had the longest pectoral fin (mean 46.2) and Arctic charr from Lake GR had the smallest pectoral fin (mean 23.6, Table 2).

**Table 2.** Estimates (average values) and significance scores from ANOVA for residuals from 8 metric and meristic traits for all 9 populations. GR-U= number of gill rakers on the upper arch, GR-L= Number of gill rakers on the lower arch, LG= length of longest gillraker, M= Length of mandibula, L=Length of lamella, LH= head length,

| Lake | GR-U | GR-L | LG | M | L | LH | EA | LP |
| --- | --- | --- | --- | --- | --- | --- | --- | --- |
| FI | 10.23*** | 15.23*** | -0.26** | -0.66* | -0.55*** | -1.00* | -1.05 | -0.99 |
| GR | -0.37 | -0.47* | 0.40** | 1.31** | 0.62*** | 1.28* | 4.42*** | 2.13** |
| LU | -0.45* | -0.26 | -0.00 | 0.72 | 0.60*** | 0.92 | -1.48 | -0.83 |
| SN | -0.23 | 0.31 | -0.05** | -0.39 | 0.62** | 0.09 | -0.50 | -0.68 |
| SV | -1.00*** | -0.35 | 0.37** | 0.83 | 1.36*** | 2.42*** | 3.38* | 1.62* |
| SØ | 0.20 | 0.00 | 0.51*** | 1.34** | 0.68*** | 1.71** | 2.35 | 2.55*** |
| TI | -0.20 | 0.45* | 0.00 | -0.36 | -0.16 | -0.90 | -0.22 | -1.40 |
| TJ | -0.65** | -0.46* | 0.49*** | 1.30** | 0.58*** | 1.62** | 0.04 | 4.43*** |
| TO | -0.61** | -0.11 | 0.72*** | 0.55 | 0.70*** | 1.51* | 0.70 | 0.44 |

EA=size of eye area, and LP= length of pectoral fin. Source population is Lake TI. Significance levels are indicated with superscript stars. \*=0.05, \*\*=0.001, \*\*\*<0.001.

Clustering of the populations based on metric and meristic traits with the Linear Discriminant Analysis suggested a visual division into two major clusters (CCC: −3.942, Fig. 4, Table 3). Here populations TI, FI, LU and SN were clustered into one group, and populations GR, TO, SØ, SV, and TJ clustered into a separate group. The first canonical axis explained 36.8 % (eigenvalue 1.00) and the second axis explained 30.1 % (eigenvalue 0.84) of the variation giving a total explained variance of 60.9 % in the first two axes. Training of the individuals into the different populations with the Discriminant analysis (DA) based on metric and meristic traits resulted in 43.6 % misclassification (Log likelihood: 558.6, Table 4). Hence, 56.4 % of the Arctic charr were correctly classified into their own population, indicating that the traits differed significantly enough for the analysis to be able to correctly identify from which lake approximately 56,4 % of the Arctic charr belonged.

**Table 3.**
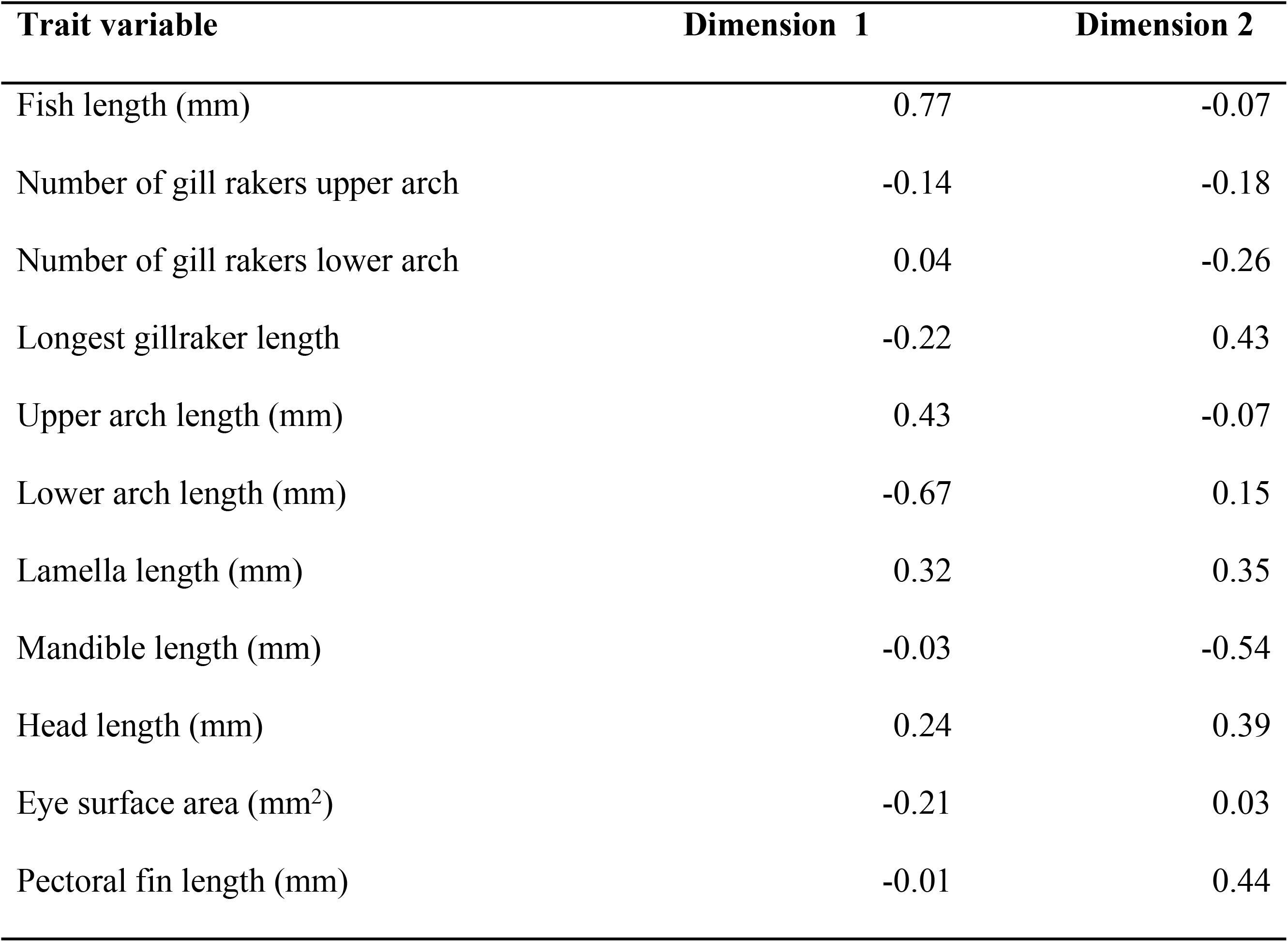
Canonical scores from discriminant analysis (DA) for residuals from 10 metric and meristic traits for all 9 populations. Canonical scores describe the coordinates for each trait in a two-dimensional system where Dimension 1 is along the X-axis and Dimension 2 is along the Y-axis (see Fig. 3).

**Table 4.**
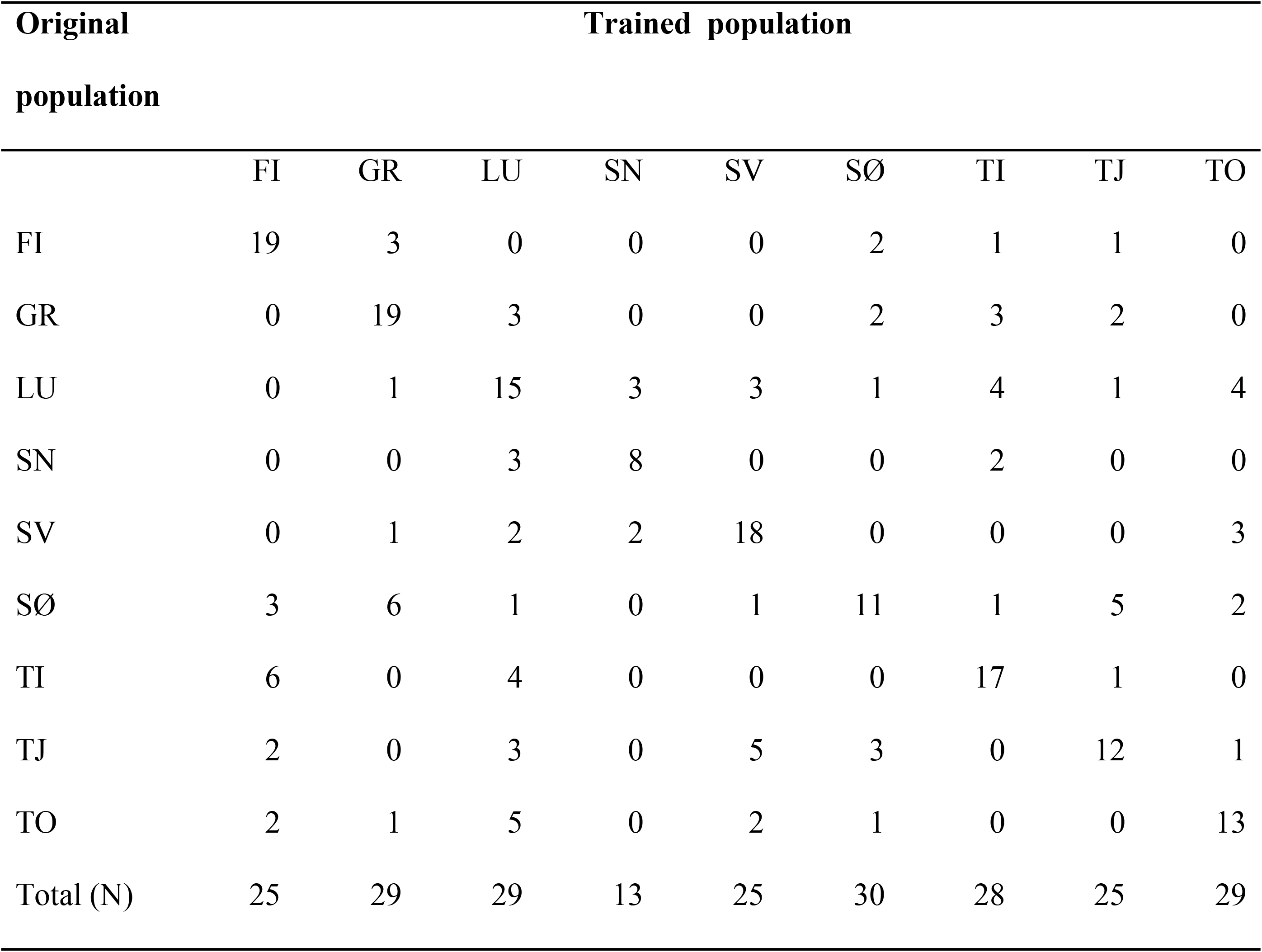
Assignment of individuals from all 9 populations back to the original populations with discriminant analysis (DA) based on residuals from 10 metric and meristic traits. Vertical lake names indicate original population, and horizontal columns indicate the number of charr individuals assigned to the different populations.

| Original<br>population | Trained population |  |  |  |  |  |  |  |  |
| --- | --- | --- | --- | --- | --- | --- | --- | --- | --- |
|  | FI | GR | LU | SN | SV | SØ | TI | TJ | TO |
| FI | 19 | 3 | 0 | 0 | 0 | 2 | 1 | 1 | 0 |
| GR | 0 | 19 | 3 | 0 | 0 | 2 | 3 | 2 | 0 |
| LU | 0 | 1 | 15 | 3 | 3 | 1 | 4 | 1 | 4 |
| SN | 0 | 0 | 3 | 8 | 0 | 0 | 2 | 0 | 0 |
| SV | 0 | 1 | 2 | 2 | 18 | 0 | 0 | 0 | 3 |
| SØ | 3 | 6 | 1 | 0 | 1 | 11 | 1 | 5 | 2 |
| TI | 6 | 0 | 4 | 0 | 0 | 0 | 17 | 1 | 0 |
| TJ | 2 | 0 | 3 | 0 | 5 | 3 | 0 | 12 | 1 |
| TO | 2 | 1 | 5 | 0 | 2 | 1 | 0 | 0 | 13 |
| Total (N) | 25 | 29 | 29 | 13 | 25 | 30 | 28 | 25 | 29 |

**Fig. 4.**
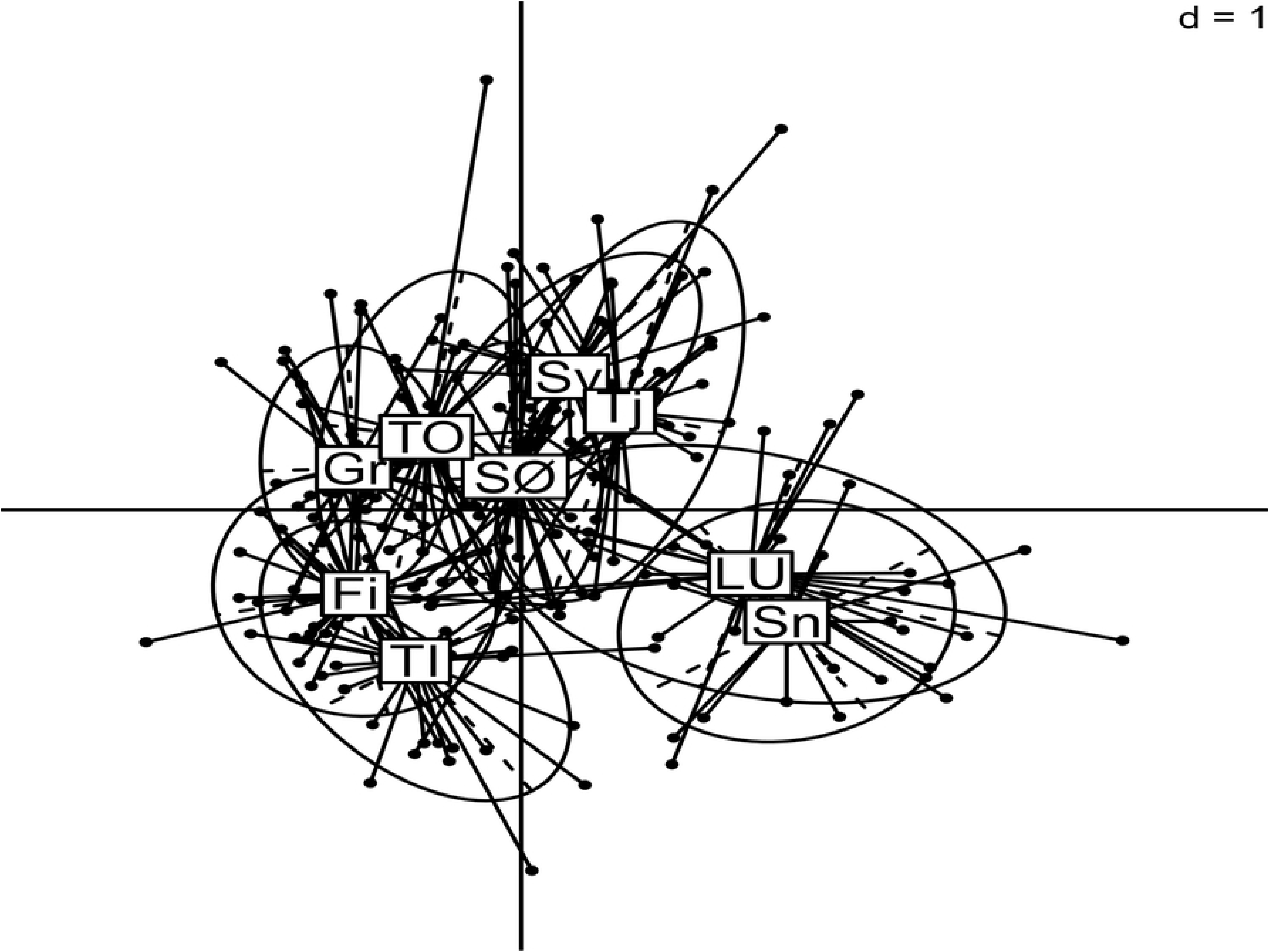
Canonical group scores and coordinates from discriminant analysis of the following morphometric characteristics for 9 lakes and 10 metric and meristic traits. Positive or negative direction of the two most influential morphometric traits is indicated along the dimensions: X – axis represents dimension 1 and Y – axis represents dimension 2.

The MANOVA analysis showed that both maximum depth and size of the lake had a significant effect (p<0.001) on the differences in morphological and meristic traits in all populations. The combined morphological and meristic traits were negatively correlated with depth (estimate: −0.95), i.e. Arctic charr in deeper lakes had for example smaller pectoral fins and fewer gill-rakers. Lake size exhibited the opposite pattern with a positive correlation (estimate: 7.71) between lake size and morphological and meristic traits (Table 5). Whether or not the lakes contained multi-or single fish species communities, significantly affected (p<0.001) the morphological and meristic traits of the populations (Table 5). The presence of several fish species in a lake was negatively correlated with morphological and meristic traits (estimate: −55.08), while single fish species communities gave a positive correlation (estimate: 122.75) with morphological and meristic traits.

**Table 5.** MANOVA table describing the effect of maximum depth, Lake size (km^2^), presence of single, or complex fish communities and individual Lakes on all morphological and meristic traits; M, LP, LH, GR-U, L, GR-L, LG, LA-L, EA, UA-L, PC1 and PC2.

|  | Df | Sum Sq | Mean Sq | F value | P-value |
| --- | --- | --- | --- | --- | --- |
| Max_depth | 1 | 74598 | 74598 | 28.15 | <0.001 |
| Lake_size | 1 | 177924 | 177924 | 67.14 | <0.001 |
| Fish_com_type | 1 | 91881 | 91881 | 34.67 | <0.001 |
| Lake name | 4 | 88007 | 22002 | 8.30 | <0.001 |

The rates of trait divergence from Lake Tinnsjøen to the translocated populations based on Darwin’s (kilodarwin’s) and Haldane’s ranged from −7.81 to 1.39, and from −0.48 to 0.20, respectively (Table 6). The three populations that exhibited the highest rates of divergence according to Darwin’s were LU, SN and TJ, with rates over 4 in most traits except for number of GR-U and GR-L (and LG for SN). All populations exhibited a negative change, i.e. a reduction in eight of the morphological traits; M, LP, LH, L, LG, LA-L, EA and UA-L. For Haldane’s, the populations with the fastest divergence rates were SØ, and TJ with positive or negative values over 0.10. The trait that has had the highest divergence across population (except for SN) was GR-L with an average negative divergence rate of −0.26 across populations, meaning that the number of GR-L has decreased in the translocated populations in comparison with the source population (Table 6).

**Table 6.** Divergence in Darwin (*10^3^) (A) - and Haldane units (B) for 10 metric and meristic traits from TI to the 8 translocated populations. GR-U= number of gill rakers on the upper arch, GR-L= Number of gill rakers on the lower arch, LG= length of longest gillraker, M= Length of mandibular, LA-L=length of lower gill arch, LA-U=length of upper gill arch, LH= head length, EA=size of eye area, and LP= length of pectoral fin. Source population is Lake TI.

| Darwin's ($\times 10^3$) | TO | SØ | FI | GR | LU | SN | SV | TJ |
| --- | --- | --- | --- | --- | --- | --- | --- | --- |
| M | 0.61 | -4.04 | 0.19 | 1.35 | -6.03 | -5.96 | -1.81 | -5.90 |
| LP | 0.33 | -3.92 | -0.17 | 0.72 | -5.12 | -5.48 | -1.97 | -5.99 |
| LH | 0.24 | -3.31 | 0.07 | 1.39 | -5.14 | -5.29 | -1.81 | -4.96 |
| GR-U | 0.42 | -0.39 | -0.19 | 0.17 | 0.26 | 0.04 | 0.84 | 0.47 |
| L | -1.14 | -5.13 | 0.01 | 0.39 | -7.23 | -7.60 | -4.57 | -6.85 |
| GR-L | 0.36 | 0.29 | 0.29 | 0.60 | 0.46 | 0.09 | 0.52 | 0.60 |
| LG | -1.35 | -3.58 | 0.33 | 0.48 | -4.27 | -3.49 | -1.83 | -4.98 |
| LA-L | -0.35 | -4.19 | -1.16 | 0.48 | -5.71 | -6.39 | -2.26 | -6.08 |
| EA | 0.87 | -3.65 | 0.34 | 0.74 | -5.08 | -5.62 | -2.06 | -5.03 |
| LA-U | -0.74 | -4.57 | -0.66 | 0.30 | -6.90 | -7.81 | -2.73 | -6.64 |

| Haldane's | TO | SØ | FI | GR | LU | SN | SV | TJ |
| --- | --- | --- | --- | --- | --- | --- | --- | --- |
| M | 0.03 | 0.10 | 0.04 | 0.09 | 0.01 | 0.02 | 0.04 | 0.10 |
| LP | 0.03 | 0.12 | 0.05 | 0.11 | 0.01 | 0.03 | 0.05 | 0.09 |
| LH | 0.03 | 0.15 | 0.05 | 0.10 | 0.01 | 0.02 | 0.06 | 0.13 |
| GR-U | -0.12 | -0.03 | -0.04 | -0.07 | 0.05 | -0.04 | 0.18 | -0.15 |
| L | 0.02 | 0.14 | 0.04 | 0.09 | 0.02 | 0.04 | 0.06 | 0.15 |
| GR-L | -0.21 | -0.38 | -0.17 | -0.48 | -0.37 | -0.06 | -0.25 | -0.20 |
| LG | -0.01 | 0.10 | 0.03 | 0.04 | 0.01 | 0.02 | 0.03 | 0.10 |
| LA-L | 0.03 | 0.12 | 0.11 | 0.05 | 0.01 | 0.01 | 0.08 | 0.15 |
| EA | 0.02 | 0.09 | 0.04 | 0.04 | 0.02 | 0.01 | 0.04 | 0.09 |
| LA-U | 0.03 | 0.10 | 0.07 | 0.06 | 0.02 | 0.02 | 0.09 | 0.20 |

### Genetic analyses

Significant genetic differentiation (*F_ST_*) was found between most of the Arctic charr from the eight translocated populations and Lake Tinnsjøen Arctic charr. The exception was non-significant differentiation between Arctic charr from Lake TI and FI, TI and LU, TI and SØ, and FI and LU (Table 7). *F_ST_*’s ranged from 0.01 between Arctic charr from Lake FI and TJ, to 0.14 between Arctic charr from SN and TO. The population pairs exhibiting the highest *F_ST_* values (above 0.11) were Arctic charr from; Lake TI and GR, SN and SV, TO and TI, SØ and TI, and between SV and TO and TI (Table 7).

**Table 7.**
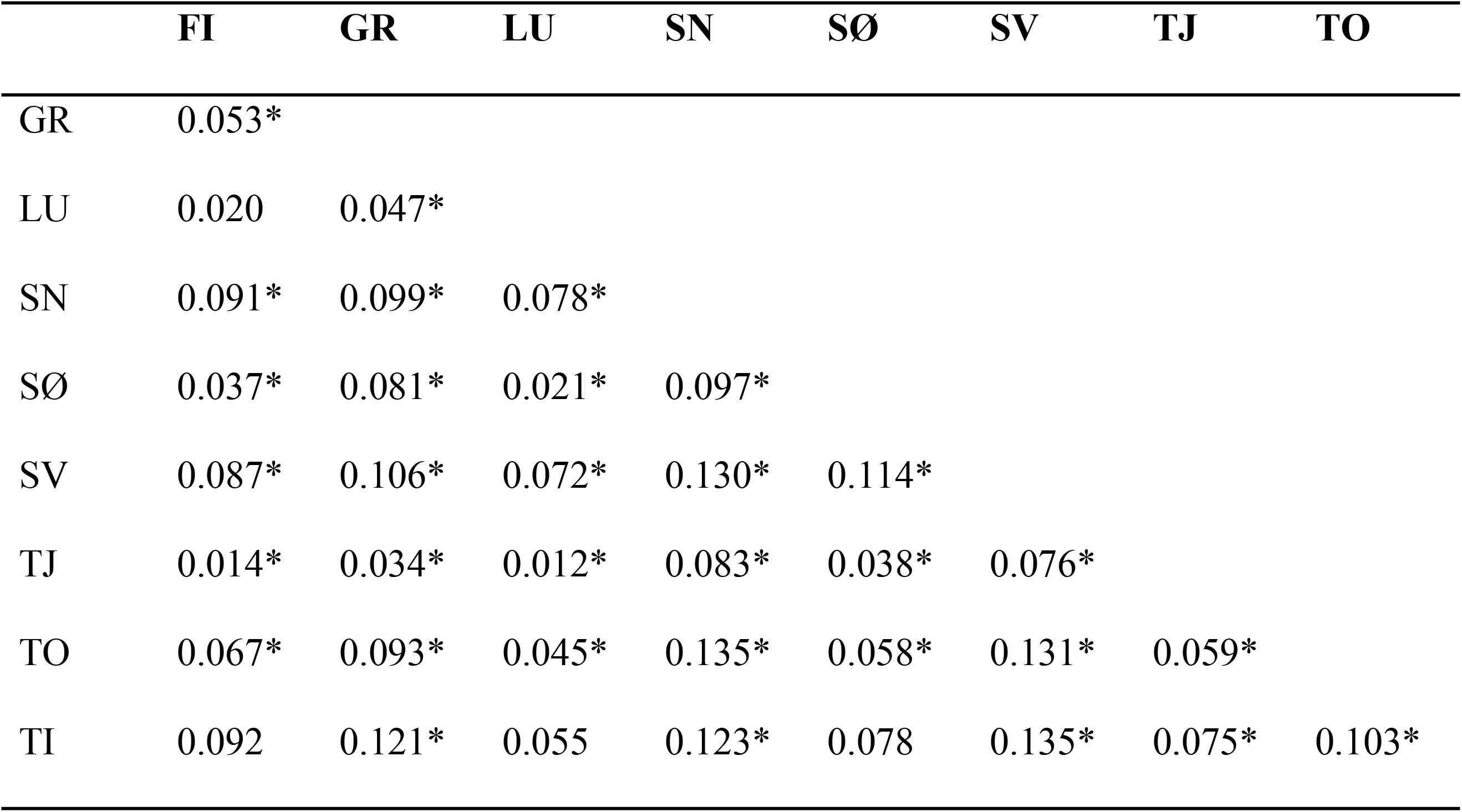
ENA-corrected F_ST_ values (pairwise) for source and translocated populations. *Significant different F_ST_ values: <0.05 *

Testing for population genetic structure using STRUCTURE, a hierarchical approach was performed where the initial run suggested 6 genetic clusters (deltaK = 5.623, mean LnP(K) = 7459.9, Fig. S2). Here, further runs were performed where the most deviating population was removed from each run to investigate population structure. The most likely partition was thus six clusters, where no significant genetic differentiation was observed between Lake TI and three of the translocated populations (FI, LU, and SØ) (Fig. 5).

**Fig. 5.**
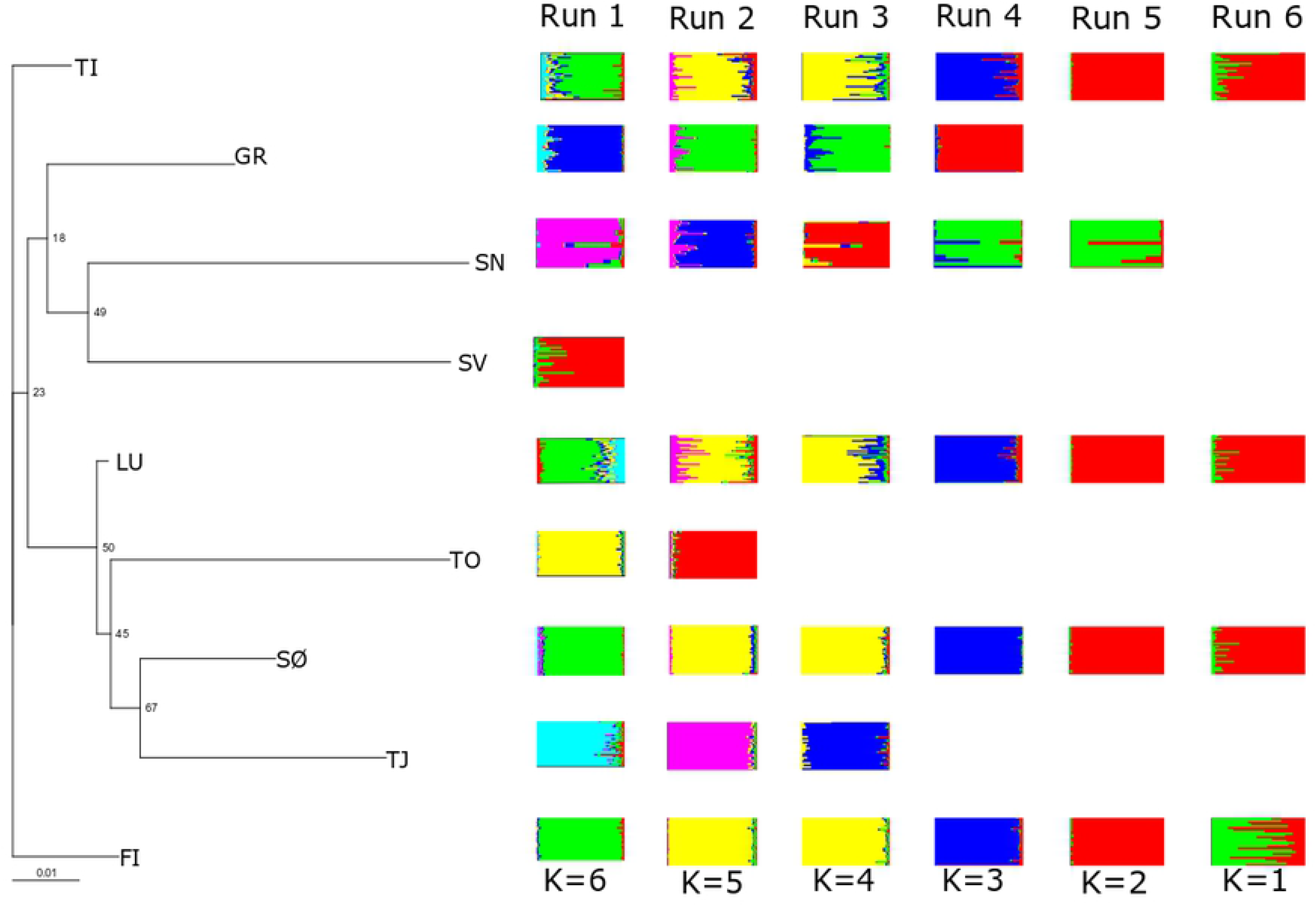
Cavalli – Sforza Neighbor joining tree (left) with branches indicating genetic relationship. STRUCTURE plot (right) where all populations are illustrated in run 1, while the most genetic differentiated population is removed singularly in each run (2 – 6). Hence run 6 consists of the most genetically similar populations.

The phylogenetic neighbor-joining tree suggested a partition into three main branches where Lake TI and FI resided on the main branch, whilst only a minor bootstrap support of 23% separated Lake GR, SN and SV populations LU, TO, SØ and TJ into the next two main branches. Minor bootstrap support (18%) separated Lake GR from Lake SN and SV, while low support (49%) separated Lake SN from Lake SV. Low bootstrap support of 50% separated Lake LU from Lake TO. Likewise, a low bootstrap value of 45 % separated Lake TO from SØ and TJ, while Lake SØ and TJ were separated by a moderate bootstrap support of 67% (Fig. 5). The second neighbor-joining tree including five outgroups showed a similar pattern where the outgroups resided in a separate branch from the remainder of the populations (Fig. S4).

### Association between genetic and phenotypic variation

The overall *F_ST_* value calculated from the genetic data was 0.072. The first 25 PCA axes that explained 50% of the total variation (Fig. 6) was used to illustrate genetic variation in the redundancy analysis. The morphological traits that best explained the genetic variation, i.e., where variation in phenotypic traits reflected genetic variation, were LG, PCA2, GR-L and L. When analyzing all morphometric traits, 6.9% of the total variation in the genetic data was associated with the phenotypic variation, and the four traits (LG, PCA2, GR-L and L) that explained the data best accounted for 3.8% (Fig. 7). With overall *F_ST_* of 0.072, the 6.9% explained variation was equivalent to an *F_ST_* of 0.005. Significance of the ordination axis was tested with an ANOVA where RDA1 was equivalent to 1.7% (p=0.001) and RDA2 to 0,9% (p=0.003) of the total variation.

**Fig. 6.**
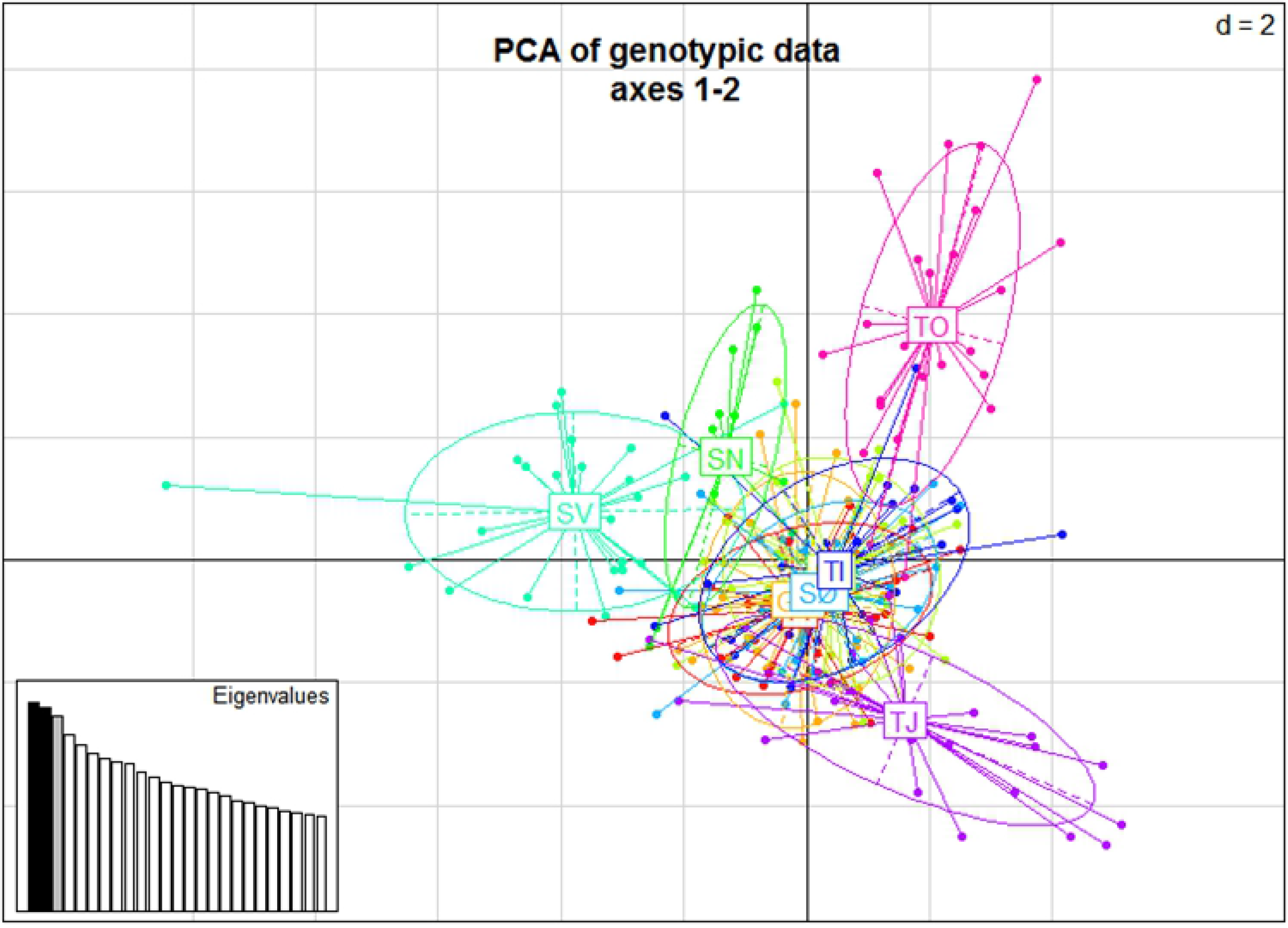
Illustration of the first two axis of principal component analysis of genetic data. PC axis one accounted 3.1% and axis two for 3.0% of the total variation.

**Fig. 7.**
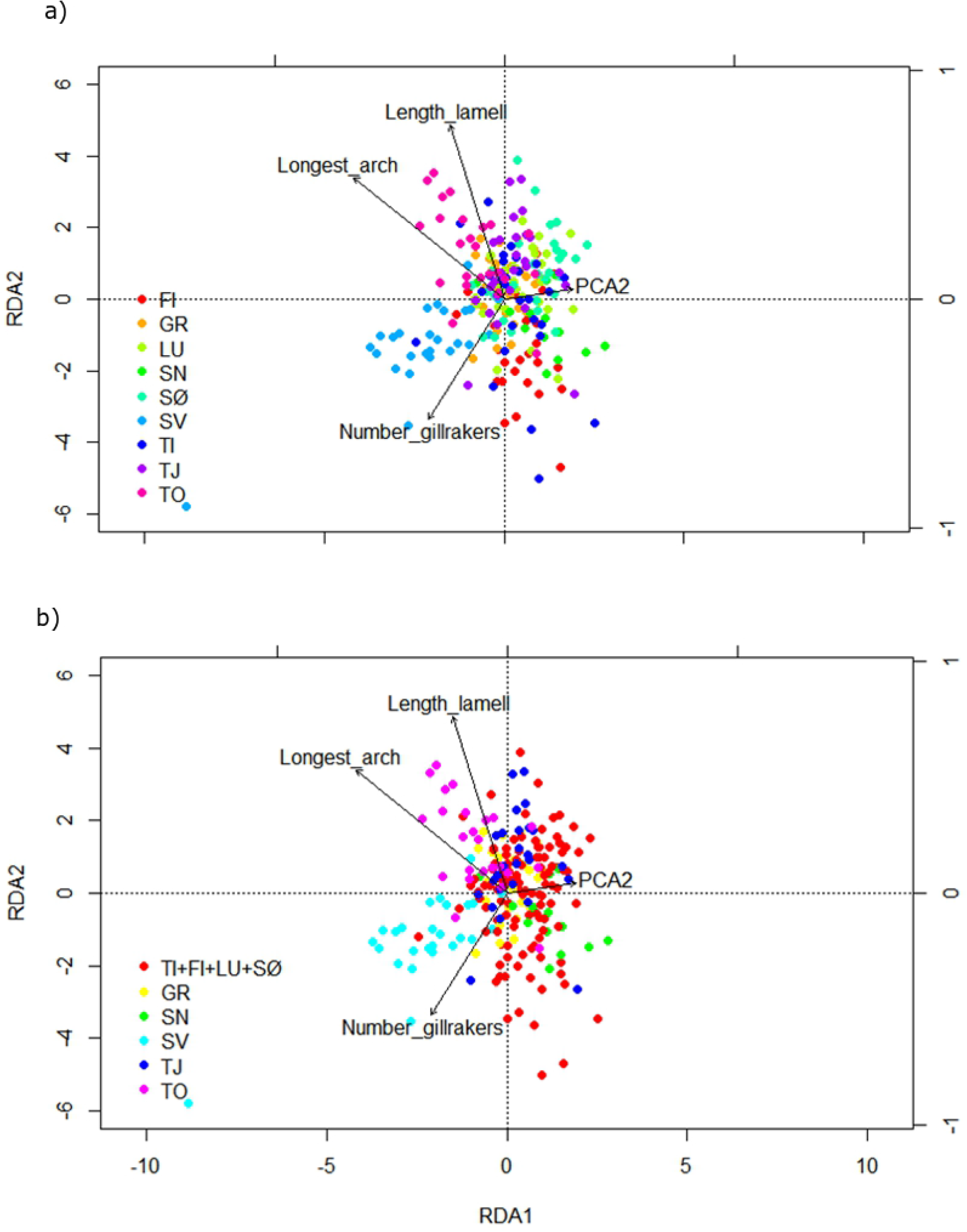
**a**. Dots represent the ordinated allele frequencies, arrows the phenotypic variables plotted as a vector. R2 = 0.038, adjusted R2 = 0.021, RDA1: 1.7% *** RDA2 0.9%**. Individuals are coded according their lake. **b.** Individuals are color coded according the six genetic clusters from the Bayesian Cluster analysis.

## Discussion

### Genetic differentiation and population structure

Significant genetic differentiation (*F_ST_*) was observed between Lake Tinnsjøen Arctic charr and five of the translocated populations, but not between Arctic charr translocated from Lake Tinnsjøen to FI, LU, SØ, and LU. This pattern was also supported by the Bayesian Cluster analysis of genetic data. Similar genetic rapid divergence has been observed for a population of translocated Arctic charr in northern Norway (78), and also in other translocated fish species such as Arctic grayling (*Thymallus thymallus*) and vendace (*Coregonus albula*) (101, 102), suggesting that northern freshwater fish species inhabit the potential to rapidly diverge in new environments.

All three genetic analyses showed that Arctic charr from five of the translocated populations have genetically diverged from the source population in Lake Tinnsjøen and that Arctic charr from LU, FI and SØ had not diverged from Lake Tinnsjøen. There was also a high degree of differentiation among most of the Arctic charr from the translocated populations as five of the populations clustered separately in the population genetic analysis, as well as significant genetic differentiation between Arctic charr from all translocated populations except from FI-LU. Previous studies of translocated fish populations have shown that genetic drift (e.g. 78, 102) is a major driver of divergence between the source and translocated populations as a translocation may be a subsampling of the allelic pool of the source population. But a translocation also introduce phenotypes adapted to the source environment, into a new environment which are deemed to induce new selection regimes (10, 103). The observed genetic divergence of Arctic charr from five populations (Lake GR, SN, SV, TO and TJ) and Arctic charr from Lake Tinnsjøen, as well as the genetic difference found between Arctic charr from the translocated populations, may thus be a result of genetic drift and/or varying selection pressures in the different lakes, perhaps coupled with varying numbers of founders in the different lakes. However, studies that investigate genome-wide divergence and how random drift and selection modulate genes and larger adaptive genomic regions for the source and translocated populations are needed to fully understand the underlying mechanisms.

### Morphological divergence and rates of differentiation

Our results show that translocating Arctic charr into new environments potentially change head, gill-raker, and pectoral fin morphometrics, while the overall body shape have remained unchanged over the 25 generations. Here, both phenotypic plasticity, or genetically based selection could act to produce an adaptive phenotype. Alternatively, the changes reveal non-adaptive variants in phenotype. The observed changes may be in accordance with the theory of ecological speciation where key trophic-morphological adaptations to new ecological niches act as one of the main drivers in divergence of new adaptive forms (16). The number of gill rakers is a key trophic trait in adaptive divergence in post-glacial fishes (65, 104, 105), and are also often used as a proxy for diversification in Arctic charr (26, 50). Head shape and pectoral fins have also been shown to adapt to trophic niches (26). The Haldane’s and Darwin’s rates of trait divergence, used for insight into evolutionary processes (106), indicated divergence in traits of the Arctic charr from source- and translocated populations. The Darwin values for all populations ranged within the expected range of 0-132 of observed values for 30 species reviewed in Kinnison and Hendry (106), while Haldane values for all populations exhibited some much higher means than the average expected after 25 generations (0.09 Haldane’s), based on the review by Kinnison and Hendry (106). Another study reported similarly high values for translocated grayling (*Thymallus thymallus*) populations in a comparable time of isolation (10). Overall, the trait that exhibited the highest divergence from expected Haldane rates was the number of gill rakers on the lower arch (GR-L). Similar results in higher than expected means of Haldane rates was found in a study on morphological traits by Michaud, Power (56) in a translocated population of Arctic charr after only 25 years of isolation, which is more rapid than several documented natural cases (106). It is possible that the charr in most of the translocated lakes exhibits a decreased number of gill-rakers as a response to a decreased pelagic area (and in some cases also a lack of interspecific competition) compared to Lake TI, and thus potentially a decreased dependency on zooplankton as the main prey group.

### Association between genetic and morphological divergence

The genetic divergence correlated with morphological divergence in the different Arctic charr populations. Despite of the individual level genotypic and phenotypic variation, Arctic charr from the different populations were mostly clustering according to their lake of origin and were characterized by phenotypic differences. Although the correlation between genetic and phenotypic variation was statistically significant, the effect in general was small. Phenotypic variation in morphology appeared to be more related to environmental and lake factors than genetic variation in each population. Variation in morphological traits is often linked to resource polymorphism, however this is not necessarily linked to a corresponding genetic differentiation (107). For selection to act on given heritable trait, it is necessary with sufficient genetic variation in the translocated populations in order to respond to selection pressures and thus to adapt to the new environments (10, 106).

### Lake variables, niches and ecological opportunity

A possible explanation for the similarity in genetic structure between Arctic charr from the source and some of the translocated populations is that the shape and size of the lakes themselves acts as a driver for maintaining a similar genetic structure to the source population. While Lake TI inhabits a high structural complexity with numerous niches, the translocated populations exhibits relatively little variation in environmental variables such as size, elevation, maximum depth, and invertebrate composition. As lake morphometry and habitat complexity have been shown to promote adaptation and variation in Arctic charr (108), marginal variation in environmental variables in the lakes where the charr was stocked might have explained lack of morphological divergence amongst and within the translocated populations. However, evidence for polymorphic Arctic charr populations in similar lake types has been found. For example, in numerous lakes in Russia and Greenland, sympatric morphs of Arctic charr are found at a broad specter of altitudes, some lakes also corresponding in size of the lakes in this study (27, 108). However, these Arctic charr populations have been present in their respective lakes since their ancestors migrated into the lakes after the last glaciation. In contrast, Loch (109) found divergent morphological traits after only two generations in a stocking experiment of *Coregonus clupeaformis*. In this study, however, the lakes subjected to stocking varied extensively in physio-chemical traits. Perhaps that in the absence of a strong selection pressure, 100 years or 25 generations of isolation from the source population as in this study, is a to short time frame for pronounced divergence to occur?

With a few exceptions, such as in the small and shallow Lake Vatnshlidarvatn in Iceland (110), deep lakes with few-species communities, are considered to be the most optimal systems for the formation of Arctic charr polymorphism, such as the deep Lake Tinnsjøen and Thingvallavatn (16, 26, 64). Temperature affects all biotic factors in the lake ecosystems, (111–113), and deeper and larger lakes are expected to have a slower temperature change, and a more pronounced layering of the water masses than shallower lakes (114). The diversity of zones in the lakes gives opportunity for specialized adaptations to habitats and prey items, where high intraspecific competition may select for specializations (reviewed in 23). In our study, maximum depth of the lakes seems to have had a negative effect on the variation in morphology and meristic traits of the translocated Arctic charr. Lake depth and size as well as productivity was also found as the main determining factors for recent diversification in European whitefish in postglacial lakes of northern Fennoscandia (19). In the larger (and relatively deep) monomorphic Arctic charr lakes like TO and SØ, interspecific competitors potentially reduce the amount of available niches, and thus alleviate the selection pressure for specialization to diverse niches. It is therefore likely that it is a combination of factors such as, origin, lake size, depth, strength of intra- and/or interspecific competition, and temperature that promotes the formation of variable Arctic charr populations.

### Management implications

Over half of the translocated Arctic charr populations exhibited genetic and morphological divergence from its source population after only 25 generations, supporting Arctic charr’s ability to rapidly diverge in new environments. Alternatively, a non-representative subset of offspring may have been stocked – being deviant from the mean population of its original source where the parents were taken from. There was also a correlation between the genetic and morphological changes in the translocated Arctic charr. However, it seems likely that for these changes to occur, the environment in which the Arctic charr is translocated to, need to inhabit certain characteristics, such as high niche heterogeneity in the form of large, and/or deep lakes and with a species composition that enables divergence to different niches (16, 26, 64). One important aspect of this study is that if the purpose is to study rapid divergence and adaptations in Arctic charr populations, it could be useful to select study sites in advance by first examining the lake morphometry and species composition to select the most likely lakes for presence of rapid divergence before expending multiple time-, and monetary resources in extensive survey fishing studies. By using for example depth maps of possible study lakes it is possible to pre-select the lakes that are most likely to inhabit a highly divergent Arctic charr population by studying the potential for divergence in niche use. This could also be useful with regard to management and protection, where these criteria may allow a knowledge base for determining likely polymorphic Arctic charr populations that may require specific management actions. Also, investigating other translocations to similar environments would lend more information on the stability of Arctic charr morphs.

## Acknowledgements

We would like to thank all the contributors; K. Syrstad, H. Luraas, G.Ø. Tveito, G. Meland, T.M. Tischbein, B. Grønstein, O-S. Hjelljord, K.H. Beckmann, I.M. Nymoen, Ø. Apalen, K. Luraas, H. Hegna, S.I. Bjørnerud, H.O. Karlsrud, K. Karlsrud, Å.K. Tollheim, I. Alseth, E. Eggerud, I. Stegarud, S. Gunleiksrud, R.O Dybdal, S. Garden, H. Espeland, G.O. Gøystdal, F. Miland, E. Miland, H.Ø. Husevold, A. Hovland, A. Tefre, T.F. Bolkesjø, M. Jore, J. Jore, S.J. Mollan, Ø. Kleiv, H. Kaasa, T.A. Folseraas, K. Sjøsåsen, T.A. Hvalen, A. Skeie, N. Kvåle, H. Lilleland, K. Holta, O. Holta, T. Flatin, B.H. Johansen, F. Lindeman, O.K. Eiken, G. Hovland, H.O. Øverby, D. Jaren, T. Granlien, H. Ålykkja, N. Singh, J. Kosir, OVF, Hedmark Fylkeskommune, and Tinn JFF. A special thanks to S. Andersen, and E. Robertsen for their help during fieldwork. T. L. Hanebrekke is thanked for assistance with the genetic analyses. L.K. Hagenlund is thanked for her valuable help with graphical design. Thanks to Inland Norway University of Applied Sciences and UiT the Arctic University of Norway for contributing with financial support to the study to KØ and KP. The publication charges for this article have been funded by a grant from the publication fund of UiT The Arctic University of Norway. And last but not least, E. Østbye: The project would not have been possible to do without you.

## Supporting information

**S1 Fig. Eigenvalues.** Principal component analysis (landmark – shape) of Arctic charr from the 9 populations showing percentage of variance explained by each PCA – axis.

**S2 Fig. DeltaK and mean LnP(K) values from Structure Harvester.** DeltaK and mean LnP(K) values from Structure Harvester. DeltaK values and right: LnP(K) values indicating most likely number of clusters of the 9 Arctic charr populations.

**S3 Fig. Body shape clustering.** Morphological variation from source- and translocated Arctic charr populations. Each Arctic charr individual is represented as a blue circle. The ellipse illustrates the number of clusters found (1) amongst principal component axis 1 (PC1) and principal component axis 2 (PC2). Principal component axis 3 (PC3) is not shown because of less explanatory power.

**S4 Fig. Phylogenetic neighbor-joining tree with outgroups.** Phylogenetic neighbor-joining tree of the source-(TI) and translocated populations (FI, GR, LU, SN, SØ, SV, TJ and TO), including 5 Arctic charr outgroups from other locations (BI, FE, LOR, TY and VA).

**S1 Table. Locus details**. Mean and standard error (SE) over loci for each of the 9 Arctic charr population; sample size (N), number of alleles (Na), number of effective alleles (Ne), observed heterozygosity (Ho), expected (He) and unbiased expected heterozygosity (uHe).

**S2 Table. Allelic richness.** Standardized- and private allelic richness for the 9 Arctic charr populations.

**S3 Table. Potential prey items.** Invertebrate and zooplankton caught in 8 of the 9 populations. Invertebrate and zooplankton were not sampled in Lake FI.

**S1 Data.** Genotype and phenotype frequencies, and lake information from this study.

**S1 Scripts.** All used R scripts and their explanation.

